# Enhancing the annotation of small ORF-altering variants using MORFEE: introducing MORFEEdb, a comprehensive catalog of SNVs affecting upstream ORFs in human 5’UTRs

**DOI:** 10.1101/2024.10.07.616631

**Authors:** Caroline Meguerditchian, David Baux, Thomas E Ludwig, Emmanuelle Genin, David-Alexandre Trégouët, Omar Soukarieh

## Abstract

Non-canonical small Open Reading Frames (sORFs) are among main regulators of gene expression. The most studied ones are upstream ORFs (upORFs) located in the 5’UTR of coding genes. Internal ORFs (intORFs) in the coding sequence and downstream ORFs (dORFs) in the 3’UTR have received less attention. Different bioinformatics tools permit to predict single nucleotide variants (SNVs) altering upORFs, mainly those creating AUGs or deleting stop codons, but no tool predict variants altering non-canonical translation initiation sites and those altering intORFs or dORFs.

We propose an upgrade of our MORFEE bioinformatics tool to identify SNVs that may alter all types of sORFs in coding transcripts from a VCF file. Moreover, we generate an exhaustive catalog, named MORFEEdb, reporting all possible SNVs altering existing upORFs or creating new ones in human transcripts and provide an R script for visualizing the results. MORFEEdb has been implemented in the public platform Mobidetails. Finally, the annotation of ClinVar variants with MORFEE reveals that more than 45% of UTR-SNVs can alter upORFs or dORFs.

In conclusion, MORFEE and MORFEEdb have the potential to improve the molecular diagnosis of rare human diseases and to facilitate the identification of functional variants from genome-wide association studies of complex traits.

## Introduction

Non-canonical small Open Reading Frames (sORFs) located in untranslated regions (UTR) surrounding the coding sequence (CDS, main ORF) of messenger RNAs are main regulators of translation and can themselves be translated (1–3). Upstream ORFs (upORFs) located in the 5’UTR have been the most studied sORFs for their role in translation initiation. The number of disease-causing variants creating new upORFs or altering existing ones has increased in the last years (4–6). Different kinds of variants have been identified in a wide range of rare genetic diseases, most of which are associated with decreased protein levels characterizing them as loss of function (LOF) mutations. These variants can (i) create new upstream translation initiation site (uTIS) (4, 5, 7), (ii) delete upstream stop codons (uStop) (5), and/or (iii) create new upstream stop codons (6, 8). The presence of downstream ORFs (dORFs) in the 3’UTR of mRNA as regulators of the CDS translation has also been described (9, 10). Even though less studied than upORF-variants, variants creating or altering dORFs have also been associated with rare human diseases (11). Different bioinformatics tools have been developed to annotate UTR variants from high throughput sequencing (HTS) data, primarily aimed at aiding the molecular diagnosis of rare genetic diseases (5, 12–14). These tools focus on the annotation of 5’UTR variants creating or disrupting upORFs (5, 12, 14) and/or 3’UTR variants altering different regulatory elements (i.e., polyadenylation sites) (13, 14). Despite significant efforts to study non-coding variants in human diseases (15–18), no bioinformatics tools was specifically designed to annotate variants that can create or disrupt dORFs or those that could create near-cognate TIS. Near-cognate TIS, also referred to as non-canonical TIS, are those differing by one nucleotide from the AUG codon. However, different studies have demonstrated that translation can be initiated with near-cognate TIS, with CUG being the most efficient one (i.e., the closest translation efficiency to AUG) (3, 19–23). There is, moreover, increasing evidence for the implication of variants creating near-cognate TIS in human diseases (6, 24). In addition to upORFs and dORFs, ribosome profiling analyses have identified small ORFs located within the coding sequence (internal ORFs, intORFs) but their role in the translation regulation and implication in diseases are still unknown (25).

Creation or disruption of non-canonical ORF in rare diseases have not only been associated with rare constitutional variants, but also with somatic and *de novo* variants (16, 17, 26). Moreover, several examples of common variants altering non canonical ORFs and associated with complex traits have been reported (27–29). Altogether, these observations illustrate the necessity of revealing DNA variants that could be able to create or to disrupt small ORFs along a given transcript and to characterize them in order to help understanding gene regulation in rare and common diseases.

We here updated our bioinformatics tool MORFEE (30) to predict single nucleotide variants (SNVs) that could create or disrupt non-canonical ORFs along a given transcript. In its initial version, MORFEE was used to identify pathogenic uAUG-creating variants in the 5’UTR of *PROS1* and *ENG* in protein S deficiency and hereditary hemorrhagic telangiectasia (HHT), respectively (7, 31). It is here extended to predict the creation of non-canonical TIS and the creation of non-canonical ORFs in 5’UTRs but also in the CDS and the 3’UTR of human transcripts. Besides, we applied this updated version on all possible SNVs in 5’UTRs resulting from an *in silico* mutational saturation of all human transcripts to build the MORFEEdb database cataloguing all SNVs that could alter upORFs. MORFEE was also deployed on different sets of variants from ClinVar database and from massive parallel reporter assay (MPRA) data.

## Materials and Methods

### MORFEE workflow

MORFEE is written in R and can run on all operating systems that have an R interpreter (including Linux, macOS and Windows). First, MORFEE uses ANNOVAR (32) to annotate minimal VCF files (i.e., with at least the *chr*, *position*, *reference allele* and *alternate allele* fields) containing SNVs in order to obtain needed information about the variants (i.e., gene, transcript, nomenclature). Then, the sequence of all transcripts is downloaded from GENCODE (GRCh37 or GRCh38) and variants are set on corresponding transcripts. At this step, MORFEE extracts all TIS and stop codons existing on transcripts in absence or in presence of the annotated variants and compares the information between both sequences to identify (i) newly created TIS; (ii) newly created stop, and; (iii) deleted stop codons, along a given transcript (Figure 1a). It is worth noting that alterations of AUGs are also annotated by MORFEE, and are noticed as “canonical_to_non_canonical_TIS”. Finally, MORFEE provides specific characteristics about sORF type, size, position on the transcript, Kozak sequence (together with its strength and KSS scores (24)), between others. MORFEE annotations, as detailed in the readme available on https://github.com/CarolineMeg/MORFEE, are assembled in an excel file.

**Figure 1.**
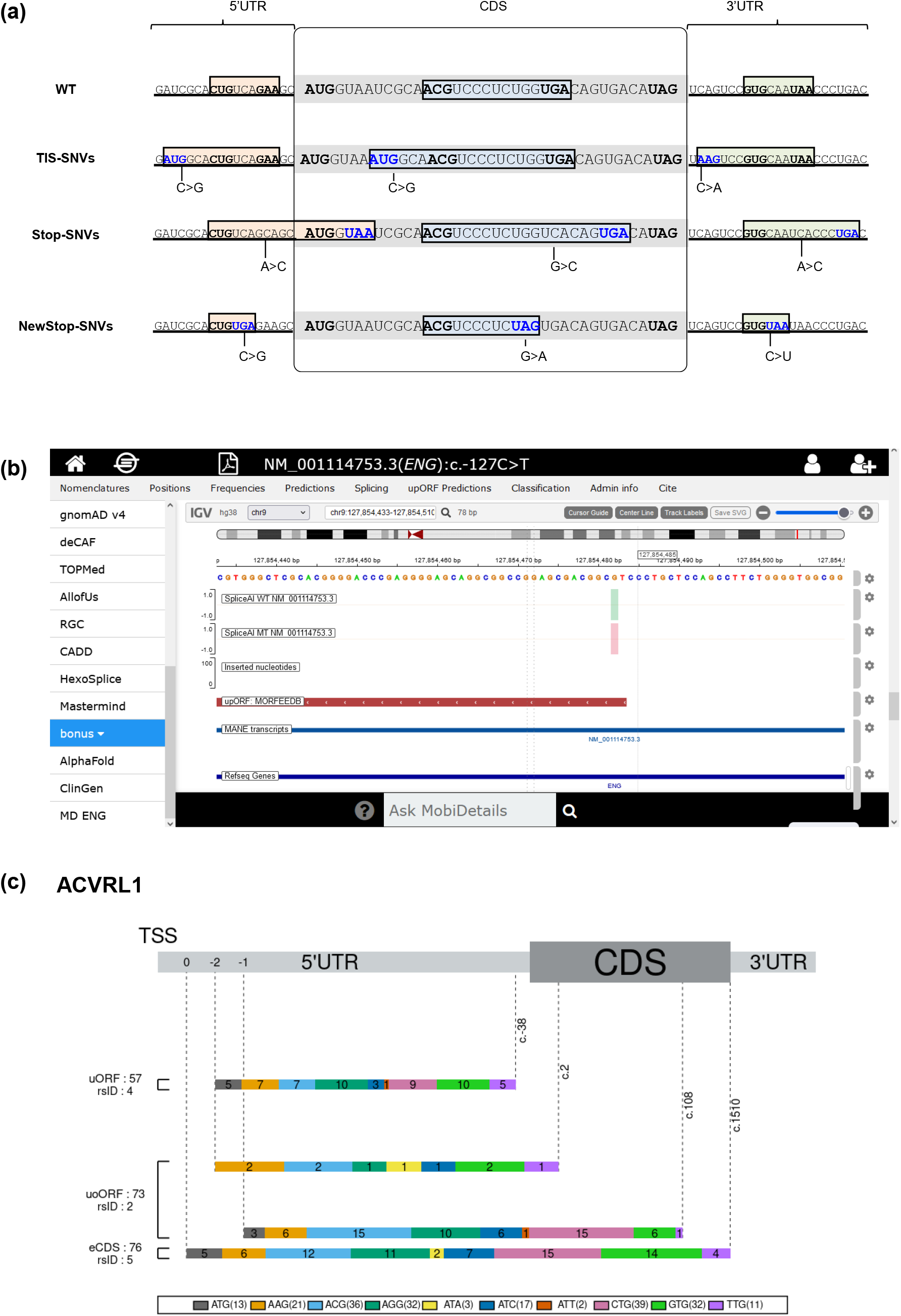
Description of MORFEE tool and its associated database MORFEEdb. (a) Annotations of single nucleotide variants (SNVs) dedicated to small open reading frames (sORFs) along a given human transcript available in MORFEE tool. The first line corresponds to a wild-type (WT) transcript sequence containing sORFs in the 5’UTR, the coding sequence (CDS) and the 3’UTR. MORFEE annotates SNVs that could create new translation initiation sites (TIS-SNVs), delete stop codons (Stop-SNVs), or create new stop codons (NewStop-SNVs). This kind of variants can create new sORFs or modify existing ones. (b) MORFEEdb output as illustrated in the public database MobiDetails. The example of c.-127C>T variant located in the 5’UTR of ENG is shown. This variant is predicted to create an upstream AUG resulting in an overlapping upstream ORF (upORF) on the MANE transcript as shown on the red dash. A table containing detailed information about the created uAUG and upORF can also be found on MobiDetails. (c) Illustration of upORFs resulting from all possible uTIS-SNVs on a given transcript extracted from MORFEEdb. A specific script allowing obtaining this graphic presentation for all human transcripts is available on https://github.com/CarolineMeg/MORFEE. The shown example corresponds to the MANE transcript (ENST00000388922.9/NM_000020.3) of *ACVRL1* gene. The total number of annotated upORFs and those resulting from variants reported in dbSNP database (https://www.ncbi.nlm.nih.gov/snp/) are indicated. The frame and position of stop codons associated with the different upORFs are shown. TSS, transcription start site; uORF, fully upstream ORF; uoORF, upstream overlapping ORF; eCDS, elongated CDS.

### Annotation of all possible upORF-SNVs in the the 5’UTR of human transcripts

In order to generate an exhaustive database containing all possible SNVs creating or altering upORFs (upORF-SNVs), we performed a mutational saturation of the 5’UTR of 55 304 transcripts from ∼17 000 coding genes. First, the most recent version of Ensembl database (GRCh38.p14) was downloaded by using BioMart package version 2.50.0 to extract the identity of all transcripts (ENST). The sequence of all transcripts was then extracted from Gencode (release 43, GRCh38.p13) and 5’UTR were defined based on the position of the main AUG. Finally, each position in the 5’UTR was *in silico* mutated with the 3 alternative nucleotides and the generated VCF files for all transcripts were annotated with the MORFEE workflow described above.

### Annotation of all ClinVar variants

All variants reported in ClinVar database were downloaded by September 2023. Variants were filtered to keep only SNVs in 5’UTR, coding and 3’UTR (i.e., in coding and non-coding exons; n = 1 048 573 SNVs; Supplemental Tables 1&2), which were annotated with MORFEE as described above.

### MPRA dataset

In order to get a rough estimate of the proportion of disease-associated dORF-SNVs that could alter the translation rates and/or RNA stability, we extracted all SNVs from Schuster et al., 2023 and annotated them with MORFEE (n = 14 497 variants). This study was dedicated to the analysis of somatic 3’UTR variants in prostate cancer (17).

## Results and discussion

### MORFEEdb, an exhaustive database for upORF-SNVs in the 5’UTR of coding genes

While most available tools mainly focus on upORFs initiated with canonical TIS (12, 14), there is accumulating evidence that some non-canonical TIS (at least CUG, GUG, UUG and ACG) can initiate the translation (8, 23) and be disease-associated (6, 8). We have upgraded the MORFEE tool, which was initially developed to annotate SNVs that create uAUG or delete uStop in the 5’UTR. We used MORFEE to generate the MORFEEdb database reporting all possible SNVs predicted to create upORFs (i.e., creation of canonical and non-canonical uTIS and of uStop) or to modify existing ones (i.e., uAUG and uStop deletion).

We *in silico* mutated the 5’UTR of 55 304 transcripts, among which 16 981 are considered as MANE Select in the Ensembl database GRCh38.p14 (Supplemental Table 3), from 17 281 coding genes (Table 1). We annotated the resulting file containing 41 536 234 SNVs with MORFEE and identified 18 886 702 upORFs as results of uTIS creations, uStop creations and/or uStop deletions in the totality of transcripts (Supplemental Table 4). The mean size of the annotated 5’UTR is 250 nucleotides containing 341 SNV-dependent upORFs in average. Transcripts’ identity, the size of their 5’UTR, the number of variants from the mutational saturation and MORFEE annotations are detailed in Table 1. According to MORFEE’s results, 14 338 457 upORFs could result from to the creation of a new uTIS, 2 788 881 from the creation of a new stop codon in the 5’UTR, and 4 798 264 from the deletion of a stop codon. This means that at least 2 503 387 upORFs are annotated on at least 2 isoforms and/or are generated from SNVs with multiple consequences (i.e., uTIS creation, uStop deletion, and/or new stop creation). While 1 472 632 of the uTIS-creating variants annotated with MORFEE could be at the origin of uAUGs in the annotated transcripts 13 615 818 create non-canonical uTIS. The annotated variants constitute the MORFEEdb database, henceforth available on this link (https://mobidetails.iurc.montp.inserm.fr/MD/static/resources/morfeedb/morfee.20231213.txt.gz).

MORFEEdb has been implemented in the public database Mobidetails and is represented schematically under the IGV panel and in a dedicated Table with detailed information about the associated upORF(s) (https://mobidetails.iurc.montp.inserm.fr/MD/, Figure 1b). Some characteristics of the corresponding upORF(s) such as the position and sequence of the uTIS and uStop, the nature of upORF (i.e., overlapping or not_overlapping), the strength of Kozak sequence surrounding the uTIS, among others are provided. Although these parameters are not exhaustive, they should help to predict the potential functional effect of upORF-SNVs. For example, 5’UTR SNVs creating overlapping upORFs are the most frequent comparing to those creating other types of upORFs (5–7, 31). Also, uTIS surrounded with strong Kozak sequences can more likely be translated and then alter the translation of the main ORF (6, 7, 33). While Whiffin et al. 2020 (5) have already annotated all possible uAUG-creating variants and uStop-deleting variants in human canonical gene transcripts, our work is the first to annotate all types of upORF-altering SNVs including those creating non-canonical uTIS. Moreover, a graphic presentation of all upORF resulting from uTIS-SNVs can be generated by using a specific script available on https://github.com/CarolineMeg/MORFEE. Figure 1c shows the example of upORFs in the 5’UTR of *ACVRL1*.

By making available MORFEEdb, with a visual and facilitated presentation on the popular Mobidetails platform, we contribute to improving the identification of upORF-SNVs in any gene of interest associated to human diseases.

### A high proportion of ClinVar SNVs can alter or create non-canonical ORFs

Despite increasing successes (16–18, 34, 35), the identification and characterization of non-coding variants causing rare genetic diseases or responsible of statistical associations observed in genome-wide genotyping/sequencing studies of complex traits remain challenging. Recent advances in ribosome profiling techniques and functional assays shed light in the role of disease-associated sORFs in translation and/or RNA stability (3, 35). The improved version of MORFEE now annotates all types of sORF-altering SNVs along a given coding transcript and is the first tool annotating dORF and intORF-SNVs. This aims to contribute to resolving molecular challenges in genetic diseases and to facilitating the identification of variants responsible for genetic association signals. This updated version of MORFEE has already enabled us to identify disease-causing uTIS--creating variants in the 5’UTR and 3’UTR of *TP53* (Li-Fraumeni syndrome) (36), in the 5’UTR *CDKN1C* with isoform-dependent effects (Beckwith-Wiedemann Syndrome) (37) and in the 5’UTR of *ENG* (HHT) (6). We here used MORFEE to annotate all possible sORF-SNVs among ClinVar variants.

Overall, 1 048 573 SNVs were reported in ClinVar by September 2023 in CDS and UTRs of human transcripts (Supplemental Tables 1&2). MORFEE annotated 6 576 upORF-SNVs in the 5’UTR among which 5 452 create new uTIS, 341 delete upstream stop codons, and 783 create new stop codons (Supplemental Table 5). Around 50% (3 202/6 576) of these variants are of unknown significance (VUS), reflecting the lack of information about this kind of variants in human diseases (Figure 2a). For instance, 15% (n = 495 variants, Supplemental Table 6) of the identified VUS among upORF-SNVs creates uTIS encompassed with strong Kozak sequence (i.e., when it contains a purine at position -3 and a guanine at position +4) and with High Kozak scores (KSS score from TIS-predictor > 0.64). Only 38 of the created uTIS correspond to AUG and 457 correspond to non-canonical uTIS including 87 corresponding to CUG (Figure 2b). In addition, 50 VUS are predicted to delete existing stop codons and are at the origin of overlapping upORFs. Of note, while the majority of pathogenic/likely pathogenic upORF-SNVs in ClinVar create uAUGs, the second most frequent pathogenic/likely pathogenic upORF-SNVs are those creating uCUG.

**Figure 2.**
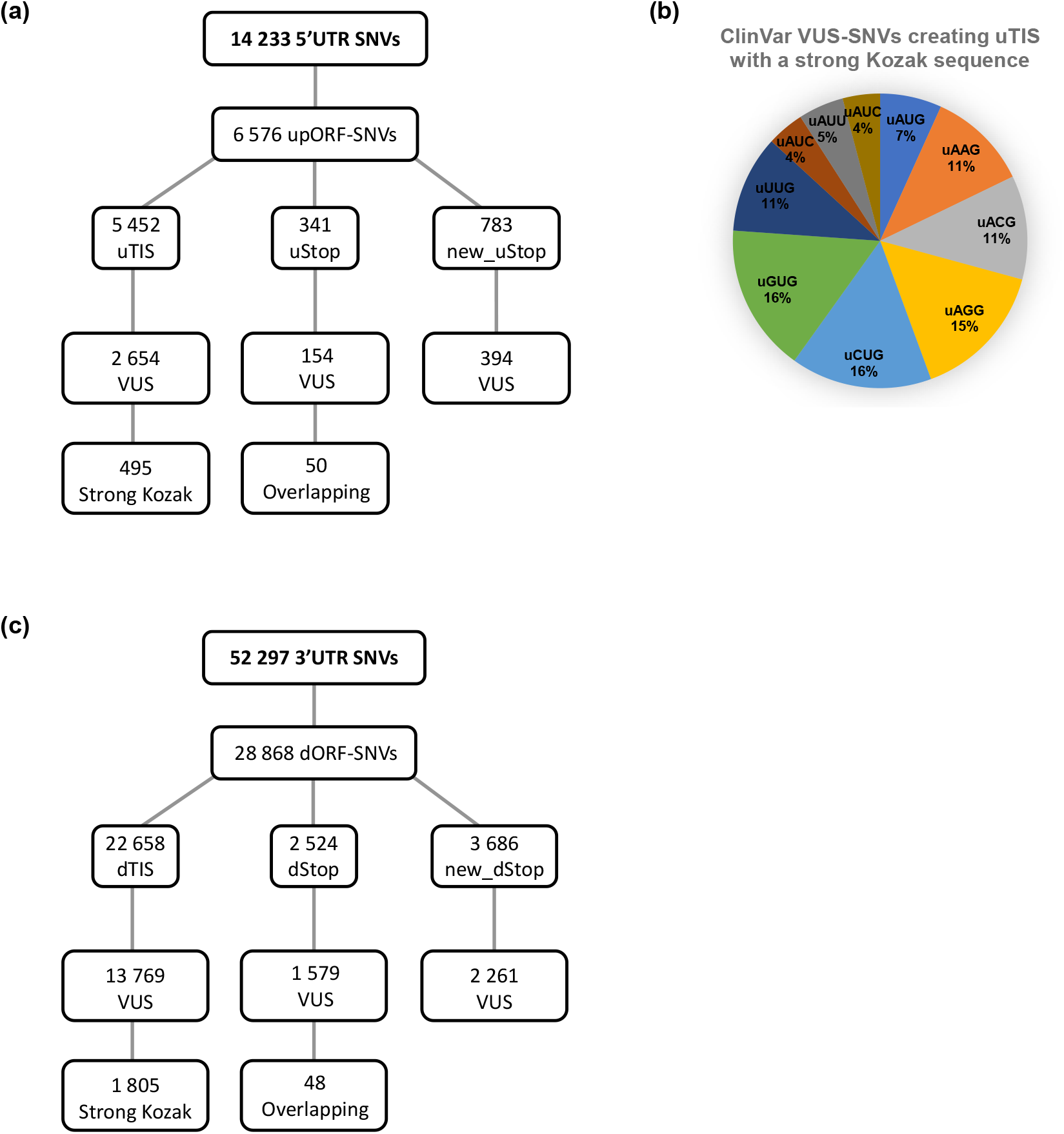
Summary of ClinVar UTR annotations with MORFEE. (a) Single nucleotide variants (SNVs) annotated with MORFEE to create or alter upstream ORF (upORFs-SNVs) among 5’UTR variants reported in ClinVar by September 2023 (n = 14 233). Variants creating new upstream translation initiation sites (uTIS), new stop codons (New_uStop), or deleting stop codons (uStop) are indicated. Variants of unknown significance (VUS) were then filtered among each category, as indicated. Strong Kozak corresponds to ClinVar SNVs classified as VUS and creating uTIS surrounded with strong Kozak sequence and a KSS score > 0.6. Overlapping corresponds to ClinVar SNVs classified as VUS and transforming fully upstream ORFs into overlapping ones by deleting stop codons. (b) Filtered 5’UTR SNVs classified as VUS and predicted to create canonical ad non canonical upstream translation initiation sites (uTIS) with strong Kozak sequence and a KSS score > 0.6. (c) Single nucleotide variants (SNVs) annotated with MORFEE to create or alter downstream ORF (dORF-SNVs) among 3’UTR variants reported in ClinVar by September 2023 (n = 52 297). Similar analysis to 5’UTR variants described in (a) was performed. dTIS, variants creating downstream translation initiation sites ; dStop, variants deleting downstream stop codons, new_dStop, variants creating new stop codons. Overlapping corresponds to ClinVar 3’UTR SNVs deleting stop codons and associated with overlapping dORFs.

In parallel, MORFEE identified 28 868 dORF-SNVs from ClinVar 3’UTR variants (Supplemental Table 7). These variants correspond to 22 658 dTIS, 2 524 dStop- and 3 686 new_dStop-SNVs among which 60% (n = 17 609 variants) are VUS with 1 805 variants creating dTIS (277 dAUG and 1 528 non-dAUGs) surrounded with strong Kozak sequences (Figure 2c).

Finally, 995 770 intORFs have been annotated with MORFEE from coding ClinVar variants (Supplemental Table 8).It is worth noting that some of the annotated variants are predicted to have multiple consequences (TIS creation, stop deletion, and/or stop creation) (Supplemental Table 9) or isoform-dependent consequences resulting from alternative 5’UTR (39). (i.e., alternative transcription start sites, alternative non-coding exons) and/or alternative main AUG (Supplemental Table 10).

On a side note, it has been shown that variants that could alter directly Kozak sequences are also able to alter translation efficiency (33). Furthermore, the presence of single nucleotide polymorphisms could also modify the access of ribosome to the 5’UTR (38). In consequence, it would be very useful to add new functions to MORFEE to annotate variants that could alter Kozak sequences of a given TIS from one side, and the potential modification of sORF-SNV effect by a nearby second sORF-SNV (i.e., haplotype effect) from the other side.

### Revisiting 3’UTR variants from MPRA data

While dORF-SNVs in the 3’UTR have been less studied comparing to upORF-SNVs, their implication in human diseases should be considered as dORFs can also regulate the translation of the main ORF. We believe that accelerated advances in high throughput function assays (i.e., MPRA techniques) will significantly clarify the implication of dORF-SNVs in human diseases.

To highlight relevant examples of disease-associated 3’UTR variants with an observed functional effect, we applied MORFEE to 14 497 somatic SNVs identified in prostate tumors and reported in Schuster et al., 2023 (17) and found 8 598 dORF-SNVs (∼60% of reported variants). The authors selected SNVs from recurrently mutated genes among the 14 497 SNVs and performed polysome-profiling-based (n = 6 892 SNVs) and RNAseq-based MPRAs to evaluate the effect of these variants on translation efficiency and RNA stability, respectively. They identified 232 3’UTR variants altering the translation efficiency and 150 variants altering the RNA stability. While the authors predicted the alteration of different regulatory elements (i.e., miRNA, polyadenylation signals and RNA binding protein motifs) by the analyzed variants, they did not take into account their potential effect on dORFs.

Very interestingly, ∼16% (37/232; dTIS = 27, dStop = 6 and New_dStop = 4) of variants altering translation and ∼49% (73/150 SNVs, dTIS = 55, dStop = 8 and New_dStop = 10) of those altering RNA stability were annotated by MORFEE as dORF-SNVs (Table 2). Whether the detected effects on translation efficiency or RNA stability are directly related to the alteration or creation of dORFs still needs to be clarified. It is worth noting that a same variant could have simultaneous consequences on different regulatory elements.

Finally, even if intORFs seems to have a low structure (2), their potential alteration with coding disease-associated SNVs still need to be explored. Of note, the annotation of all possible SNVs located within the CDS and the 3’UTR of all human transcripts with MORFEE could be necessary to expand knowledge about dORF- and intORF-SNVs, and to generate datasets of variants for functional assays. Besides, the current version of MORFEE is dedicated for the annotation of SNVs but it needs to be updated to also annotate deletions and insertions as they can create new sORFs or modify existing ones.

## Conclusion

We here propose an exhaustive catalog of all possible upORF-SNVs in human 5’UTRs and an easy-to-use tool to guide experimental analysis contributing to determine the pathogenicity of disease-associated variants. In fine, this would improve the molecular diagnosis of rare human diseases but also to identify potential common variants in complex diseases.

## Data availability

The used version of MORFEE tool is available at https://github.com/CarolineMeg/MORFEE.

SNVs resulting from the *in silico* mutational saturation of human 5’UTRs from 55 304 transcripts are available upon request.

SNVs reported in ClinVar and annotated with MORFEE to possibly create or alter internal ORFs in the coding sequence are available at 10.5281/zenodo.13872114.

## Author Contributions Statement

OS and DAT conceived the project. CM upgraded MORFEE, performed the mutational saturation and variant annotation with MORFEE. OS analyzed the data. DB TEL and EG participated to the bioinformatics development of MORFEEdb and to its implementation in Mobidetails. OS and DAT drafted the manuscript that was further shared to co-authors who read/corrected/ and approved the final manuscript.

## Acknowledgments

Statistical analyses benefited from the CBiB computing center of the University of Bordeaux. This project was carried out in the framework of the INSERM GOLD Cross-Cutting program (E.G, TE.L, D-A.T and O.S) and of the French National Research Agency (ANR) ANR-23-CE17-0042-01 program as part of the ENDOMORF project (D-A.T, O.S).

## Abbreviations

sORF: small Open Reading Frame
UTR: UnTranslated Region
CDS: CoDing Sequence
upORF: upstream Open Reading Frame
uTIS,: upstream Translation Initiation Site
uStop: upstream Stop codon
dORFs: downstream Open Reading Frame
dTIS: downstream Translation Initiation Site
dStop: downstream Stop codon
intORFs: internal Open Reading Frame
intTIS: internal Translation Initiation Site
intStop: internal Stop codon
SNV: Single Nucleotide Variant
MORFEE: Mutation on Open Reading FramE

